# Benchmarking sequence performance on the DNBSEQ-T7 using Genome in a Bottle reference genomes

**DOI:** 10.64898/2026.05.22.727100

**Authors:** Ansia van Coller, Setshaba Taukobong, Maano Malima, Samira Ghoor, Nganea Nangammbi, Enrico Roode, Martin Naicker, Victoria Cole, Brigitte Glanzmann, Craig Kinnear, Nadia Carstens

**Author notes:** These authors contributed equally (Ansia van Coller, Setshaba Taukobong).

## Abstract

Advances in sequencing technologies have improved the accuracy, throughput, and completeness of human genome characterization, enabling more reliable detection of genetic variation. Well-characterized reference genomes are critical for benchmarking sequencing platforms and bioinformatics analysis pipelines. Here, we present whole genome sequencing datasets generated for the Ashkenazi Jewish trio reference samples from the Genome in a Bottle Consortium. Libraries were prepared using three distinct MGI-based workflows: PCR-free library preparation, FastFS DNA library preparation, and Universal DNA library preparation. Sequencing was performed on the MGI DNBSEQ-T7 platform, generating a minimum of 400 million paired-end reads per sample, corresponding to 30X mean genome coverage.

Raw reads were processed using a standardized GATK bioinformatics workflow. Sequencing performance and variant detection accuracy were evaluated using the Genome in a Bottle high-confidence benchmark variant sets. All workflows demonstrated high sequencing quality and concordance with GIAB benchmark truth sets, with PCR-free libraries showing the strongest indel calling performance and lowest Mendelian violation rates across the Ashkenazi trio.

This dataset provides a resource for benchmarking DNBSEQ-T7 sequencing and bioinformatics workflows, and for evaluating the impact of library preparation strategies on whole genome variant detection performance.

## Background

The Genome in a Bottle (GIAB) Consortium provides well-characterized reference materials and benchmarking datasets for variant calling, including the Ashkenazi Jewish trio (HG002, HG003, HG004) derived from the Personal Genome Project (1). These samples are accompanied by high-confidence variant call sets generated through integration of multiple sequencing technologies, enabling robust validation of whole genome sequencing (WGS) pipelines (1,2). The Ashkenazi trio serves as a widely adopted benchmark for small variant and structural variant detection and its trio structure enables evaluation of Mendelian inheritance, providing an orthogonal approach for identifying potential errors beyond comparisons to a reference truth set (2,3).

The DNBSEQ-T7 sequencer, developed by MGI Tech, is a high-throughput platform that has been evaluated using reference genomes such as the Korean Reference Genome and NA12878 from the GIAB Consortium. These studies included both exome and whole genome sequencing, evaluating key performance metrics such as variant detection accuracy, coverage uniformity, and overall sequencing performance (4,5).

While the DNBSEQ-T7 platform has previously been evaluated using established reference genomes, including GIAB-derived standards, comparatively little attention has been given to the impact of different MGI whole genome library preparation workflows on downstream sequencing and variant calling performance. Differences in fragmentation chemistry, PCR amplification, and DNA input requirements can influence coverage characteristics, GC representation, duplication rates, and variant detection accuracy, yet these effects have not been systematically assessed across multiple DNBSEQ-T7 workflows using a standardized benchmark framework.

Here, we generated whole genome sequencing data for the GIAB Ashkenazi trio using three MGI library preparation strategies implemented under matched sequencing and analysis conditions at the SAMRC Genomics Platform. This design enables direct comparison of workflow-specific performance characteristics, including sequencing quality, coverage profiles, variant concordance with GIAB truth sets, and Mendelian consistency.

Three library preparation strategies were assessed: Universal DNA library preparation kit (fragmentation by sonication), Fast FS DNA library preparation kit (enzymatic fragmentation), and PCR-free DNA library preparation kit (enzymatic fragmentation). Sonication-based methods produce consistent fragment sizes with reduced bias, while enzymatic fragmentation offers a streamlined workflow suitable for low-input samples. PCR-free approaches reduce amplification bias and improve representation of GC-rich regions, facilitating more accurate detection of structural variants and copy number variation (6–9).

This study presents a WGS dataset for the GIAB Ashkenazi trio generated on the DNBSEQ-T7 platform across these three library preparation strategies, together with benchmarking against GIAB truth sets. The dataset provides a resource for evaluating sequencing performance, benchmarking variant calling pipelines, and guiding selection of library preparation methods based on sample quality and analytical requirements.

## Methods

### Samples

Formalin-fixed paraffin-embedded (FFPE) curls of the Genome in a Bottle (GIAB) Ashkenazi trio, HG002, HG003, and HG004, were obtained from Horizon Discovery (GM24385, GM24149, GM24143). Genomic DNA was extracted using the DNeasy Blood and Tissue Kit (Qiagen) according to the manufacturer’s instructions. DNA quality was assessed using the Qubit 4.0 Fluorometer (Thermo Fisher Scientific, Waltham, MA, USA) and the Qubit dsDNA HS Assay kit Qubit, purity assessment with NanoDrop One Spectrophotometer (Thermo Fisher Scientific, Waltham, MA, USA, and DNA integrity evaluation using the Agilent TapeStation 4200 and (Agilent, Santa Clara, CA, USA). All samples met the criteria for WGS library preparation

### Library preparation and sequencing

Whole genome sequencing libraries were prepared using three MGI workflows with reagents sourced from MGI (MGI, Shenzhen, China): (i) MGIEasy Universal DNA Library Prep Set with Covaris Focused-ultrasonicator (Covaris, Woburn, MA, USA) sonication (1000 ng input), (ii) MGIEasy PCR-Free FS Library Prep Set (500 ng input), and (iii) MGIEasy Fast FS DNA Library Prep Set (25 ng and 200 ng inputs). Single-stranded, circularized libraries were generated using the MGIEasy Circularization Kit (MGI, Shenzhen, China). Libraries were then loaded onto a flow cell using the MGIDL-T7, followed by sequencing on the DNBSEQ-T7 using paired-end 150 bp reads using the DNBSEQ-T7RS High-Throughput Sequencing Set (FCL PE150) (MGI, Shenzhen, China).

### Data Records

Raw sequencing data have been deposited in the Sequence Read Archive (SRA) under BioProject ID PRJNA1465120. The dataset comprises whole-genome sequencing data for HG002, HG003, and HG004 generated using PCR-free, Fast FS (25 ng and 200 ng), and Universal library preparation methods. The Variant call files (VCFs) for each sample has been uploaded to Zenodo (doi: 10.5281/zenodo.20340430).

### Technical Validation

Technical validation demonstrated consistently high sequencing and variant calling performance across all DNBSEQ-T7 library preparation workflows, with PCR-free libraries showing the strongest overall indel detection accuracy and Mendelian consistency.

### Sequencing quality metrics

Sequencing quality metrics were evaluated using FastQC (v0.12.1) (6) and MultiQC (v1.25.2) (7), including Q30 scores, GC content, duplication rate, insert size and read yield. All libraries demonstrated high sequencing quality (Q30 >97%) and high alignment rates (>99% to GRCh38). Summary metrics are provided in Table 1.

**Table 1.**
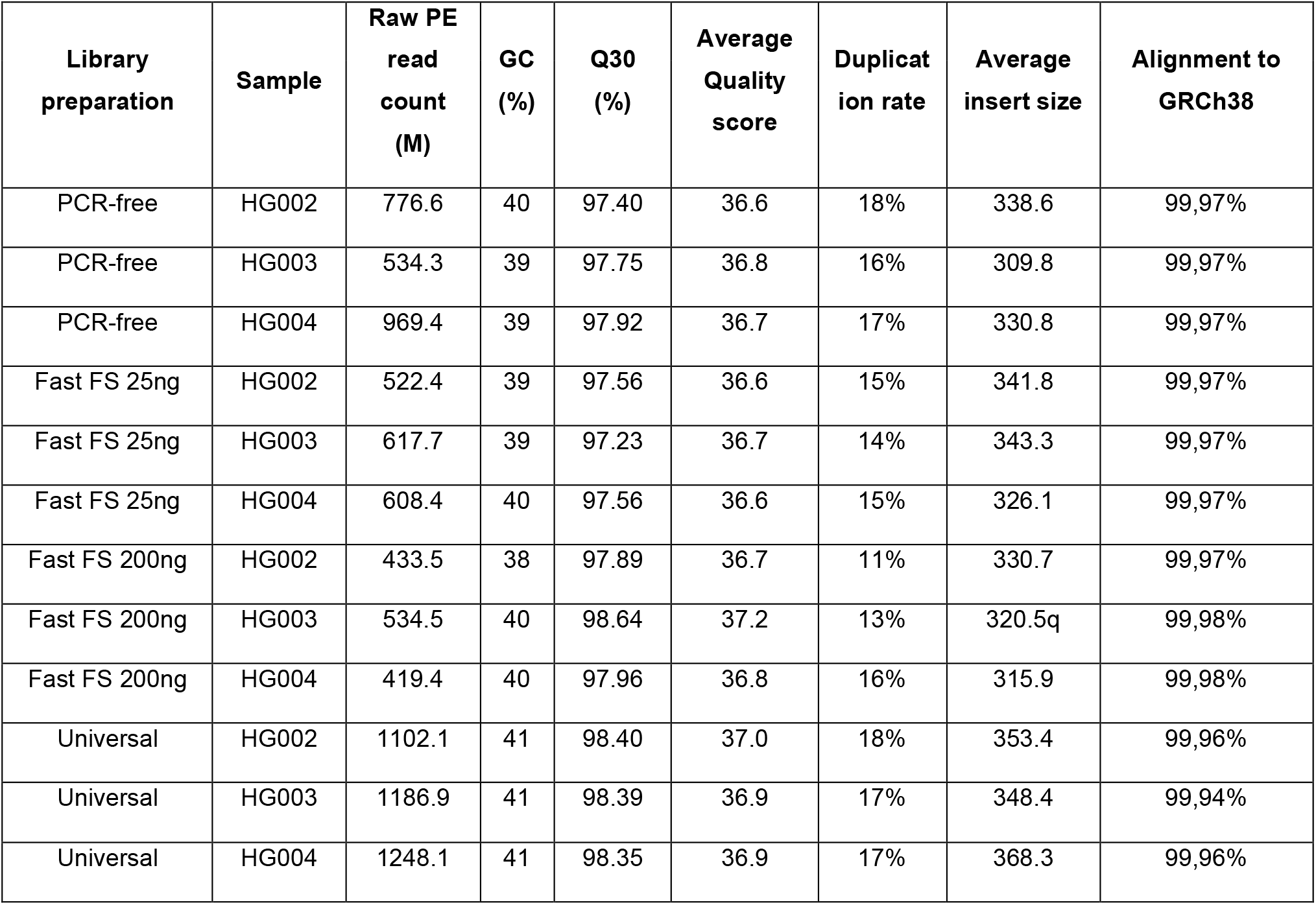

| Library preparation | Sample | Raw PE read count (M) | GC (%) | Q30 (%) | Average Quality score | Duplication rate | Average insert size | Alignment to GRCh38 |
| --- | --- | --- | --- | --- | --- | --- | --- | --- |
| PCR-free | HG002 | 776.6 | 40 | 97.40 | 36.6 | 18% | 338.6 | 99,97% |
| PCR-free | HG003 | 534.3 | 39 | 97.75 | 36.8 | 16% | 309.8 | 99,97% |
| PCR-free | HG004 | 969.4 | 39 | 97.92 | 36.7 | 17% | 330.8 | 99,97% |
| Fast FS 25ng | HG002 | 522.4 | 39 | 97.56 | 36.6 | 15% | 341.8 | 99,97% |
| Fast FS 25ng | HG003 | 617.7 | 39 | 97.23 | 36.7 | 14% | 343.3 | 99,97% |
| Fast FS 25ng | HG004 | 608.4 | 40 | 97.56 | 36.6 | 15% | 326.1 | 99,97% |
| Fast FS 200ng | HG002 | 433.5 | 38 | 97.89 | 36.7 | 11% | 330.7 | 99,97% |
| Fast FS 200ng | HG003 | 534.5 | 40 | 98.64 | 37.2 | 13% | 320.5q | 99,98% |
| Fast FS 200ng | HG004 | 419.4 | 40 | 97.96 | 36.8 | 16% | 315.9 | 99,98% |
| Universal | HG002 | 1102.1 | 41 | 98.40 | 37.0 | 18% | 353.4 | 99,96% |
| Universal | HG003 | 1186.9 | 41 | 98.39 | 36.9 | 17% | 348.4 | 99,94% |
| Universal | HG004 | 1248.1 | 41 | 98.35 | 36.9 | 17% | 368.3 | 99,96% |

### Bioinformatics Processing

Reads were trimmed using Trimmomatic (v0.40-rc1) (8). And aligned to GRCh38p14 using BWA-MEM (v0.7.18) (9). BAM files were downsampled to 400 million read pairs per sample to enable direct cross-method comparison. Processing followed Genome Analysis Toolkit (GATK) (v4.6.1) best practices (10), which including duplicate marking, base quality score recalibration (BQSR), and variant calling with HaplotypeCaller.

Coverage statistics were calculated using through GATK (v4.6.1) picardtools and mosdepth (v0.3.8) (11). Coverage profiles were consistent across methods. Coverage distributions are visualized in Figure 1, showing coverage uniformity across the library preparation methods.

**Figure 1:**
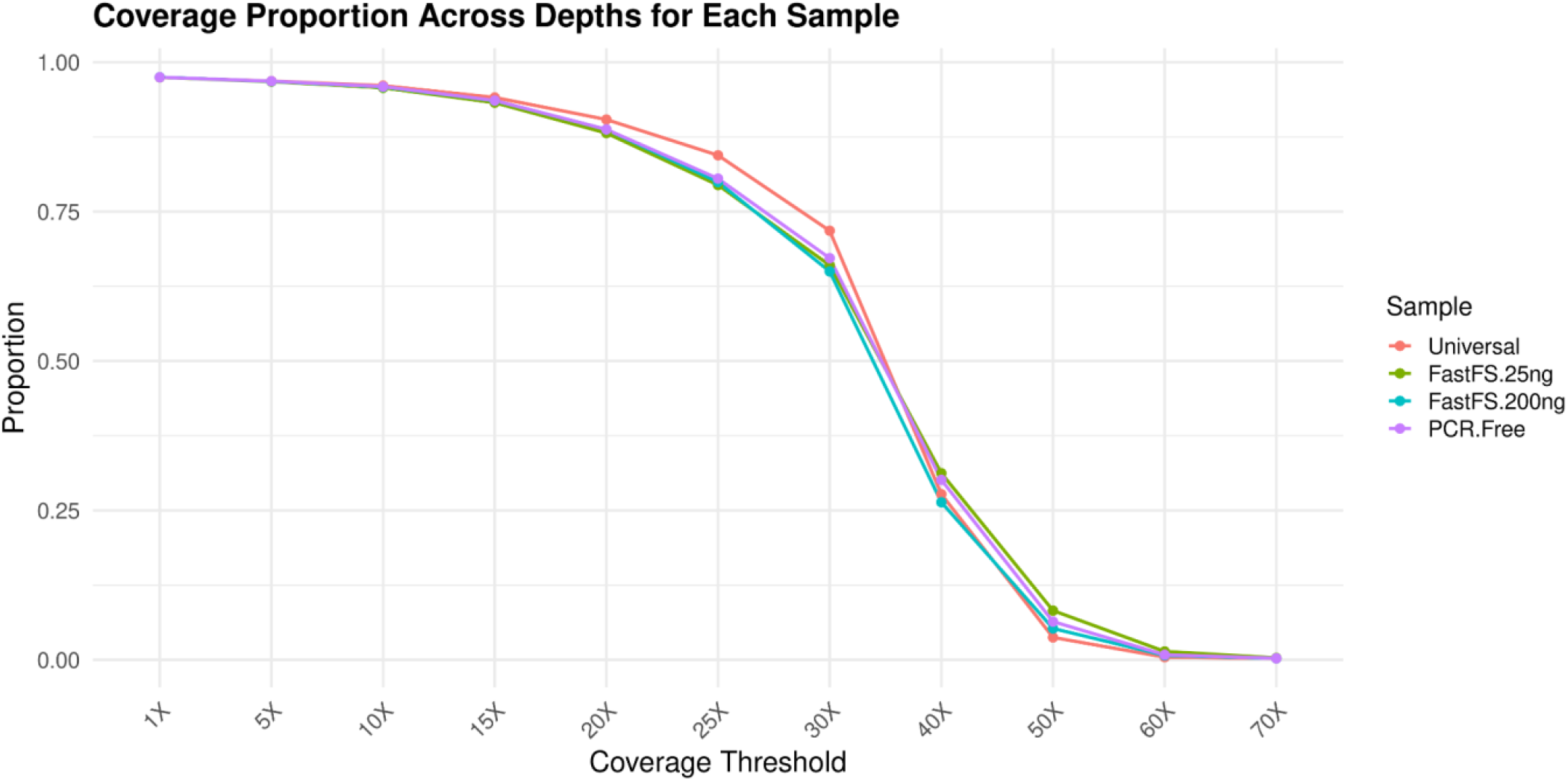
Coverage distribution for HG002 for each library preparation method sequenced on the DNBSEQ-T7. BAM files were downsampled to 400M PE reads. Results for HG003 and HG004 were comparable.

### Benchmarking against Genome in a Bottle Truth sets

Variant callsets were benchmarked against GIAB v4.2.1 high-confidence regions using hap.py (v0.3.12) (12). Performance was assessed using precision, recall, and F1 score for SNVs and indels (13). Results are summarized in Figure 2 (SNV) and Figure 3 (Indels). All methods showed high and consistent SNV performance (F1 ∼0.98–0.99).

**Figure 2:**
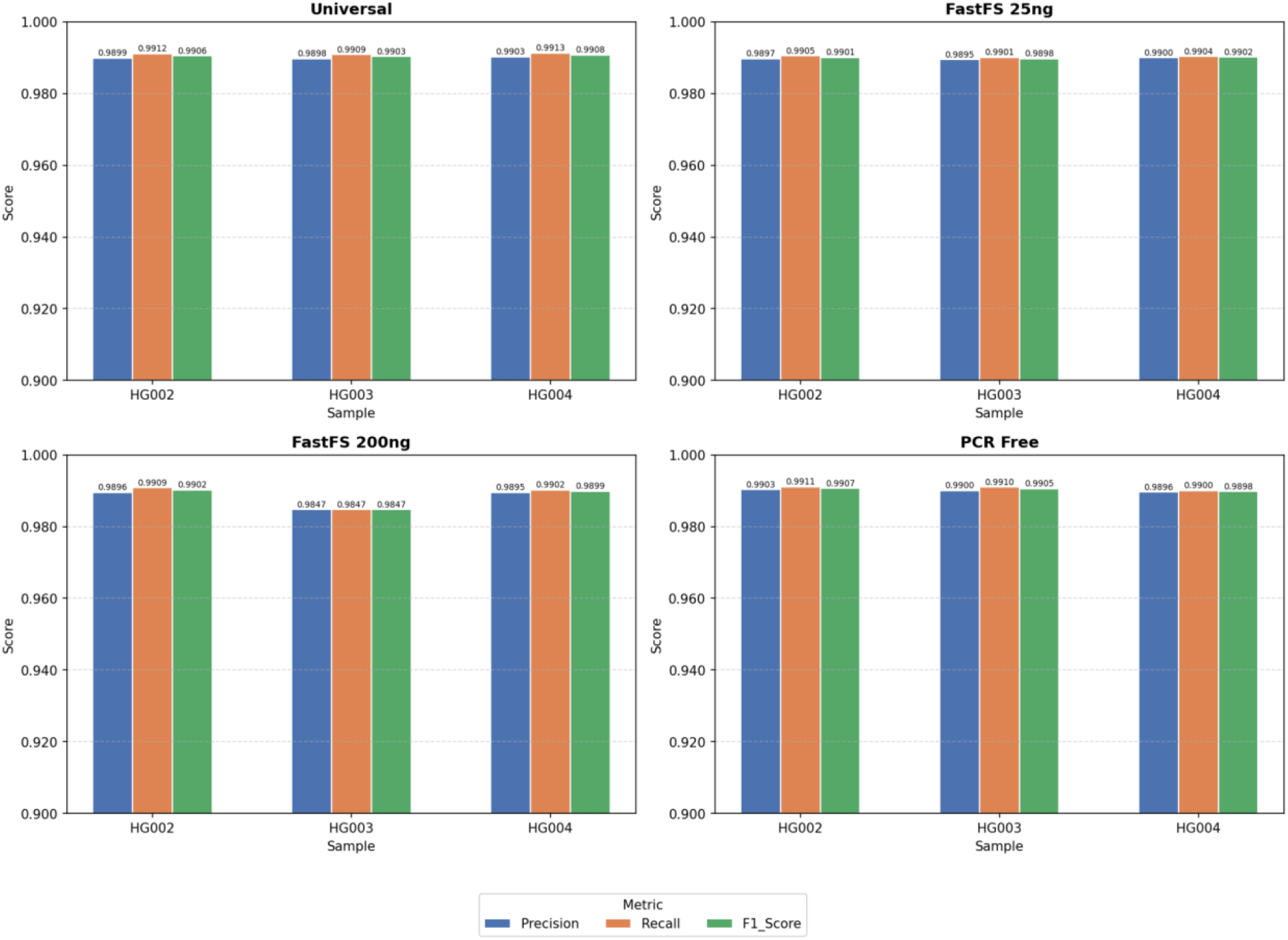
Comparison of SNV variant calling performance across the different library preparation methods (Universal, FS 25 ng, FS 200 ng, and PCR-free), shown as bar graphs of Precision, Recall, and F1-score.

**Figure 3:**
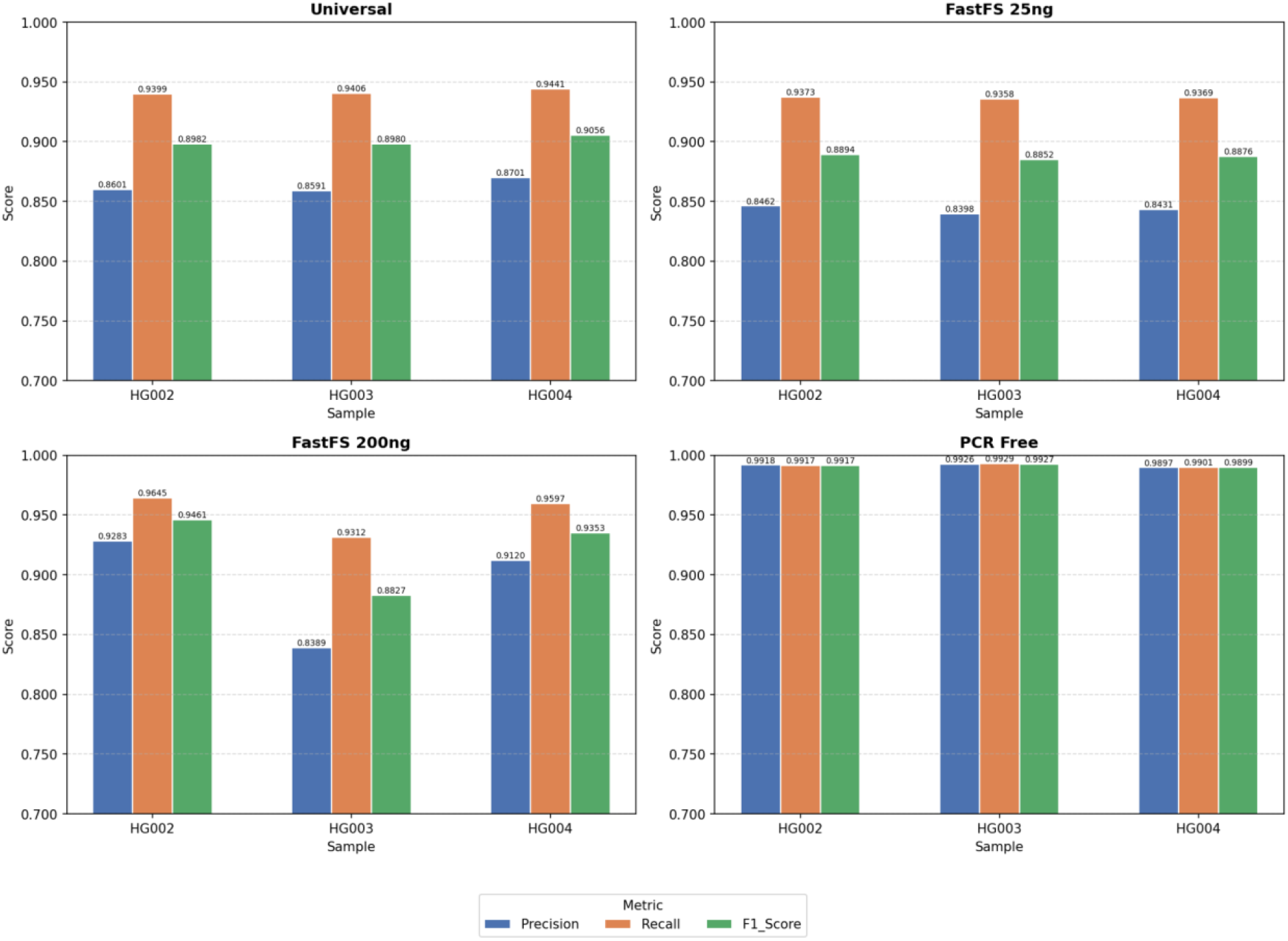
Comparison of Indels variant calling performance across the different library preparation methods (Universal, FS 25 ng, FS 200 ng, and PCR-free), shown as bar graphs of Precision, Recall, and F1-score.

Indel performance varied across methods, with PCR-free libraries achieving the highest and most balanced performance across all samples

Variant concordance across library preparation methods was evaluated by comparing overlapping and unique variant calls. Figure 4, illustrates the concordance of the different library preparation methods for HG002 for SNVs and Indels. Trio-based validation was performed using RTG Tools (v3.13) (14) to assess Mendelian consistency. Mendelian violation rates were low across all methods, with PCR-free libraries showing the lowest violation rate (Table 4).

**Table 4:** Mendelian Concordance and Violation rates for each library preparation method.

| Library preparation method | Concordance of HG002 and HG003 (%) | Concordance of HG002 and HG004 (%) | Concordance of HG002 and HG003/HG004 (%) | Records with indeterminate consistency status due to incomplete calls (%) | Violation of Mendelian constraints (%) |
| --- | --- | --- | --- | --- | --- |
| Universal | 99.99 | 99.99 | 99.41 | 0.10 | 0.59 |
| FastFS 200ng | 99.98 | 99.99 | 99.58 | 0.14 | 0.42 |
| FastFS 25ng | 99.99 | 99.99 | 99.36 | 0.11 | 0.64 |
| PCR-free | 100.00 | 99.99 | 99.86 | 0.09 | 0.14 |

**Figure 4:**
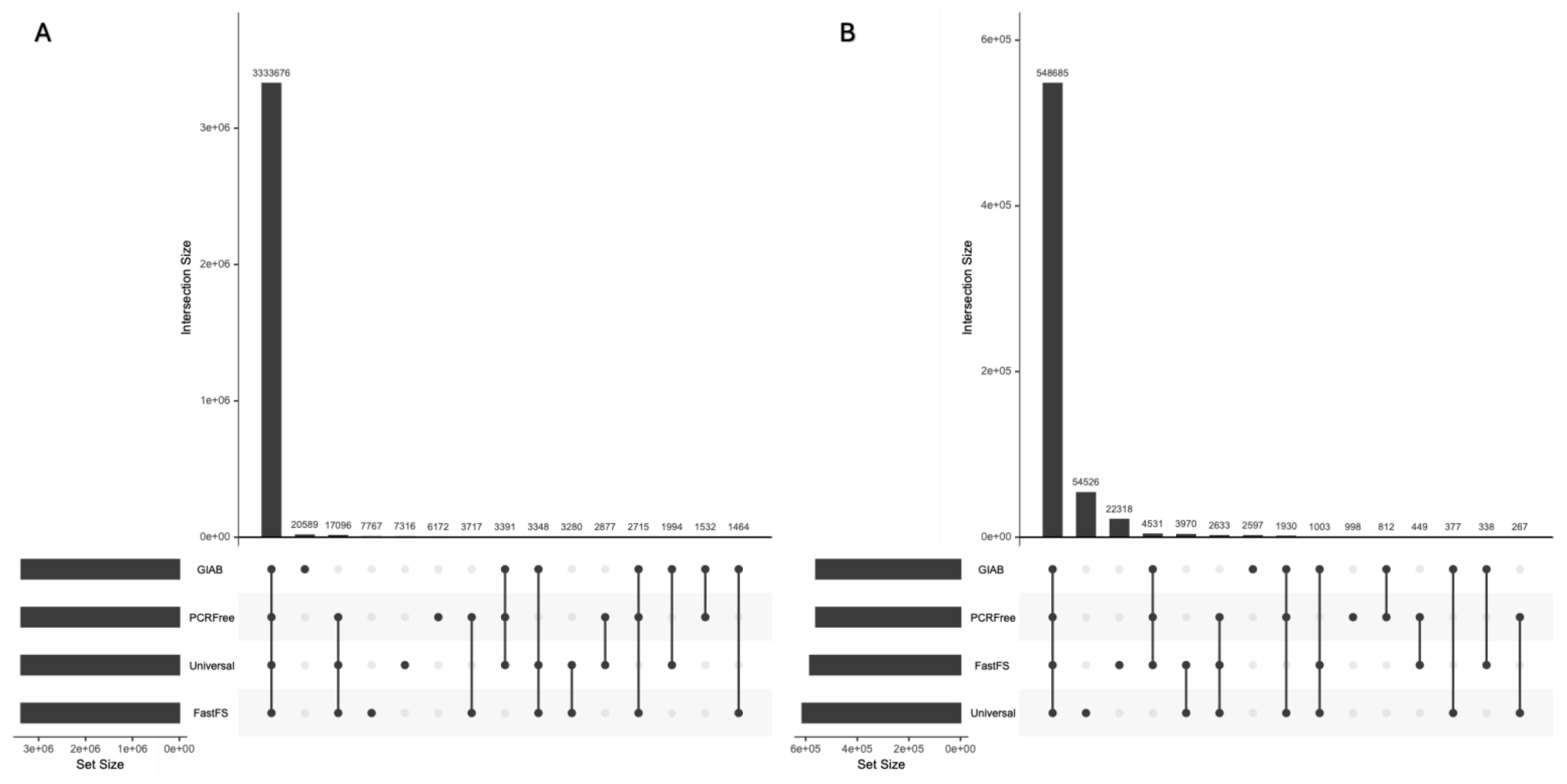
UpSet plot illustrating concordance of variant calls across library preparation methods for HG002 sequenced on the DNBSEQ-T7 for (A) SNVs and (B) Indels. HG003 and HG004 had comparable results.

## Data availability

All data generated are publicly available without restriction. Raw sequencing data have been deposited in the Sequence Read Archive (SRA) under BioProject ID PRJNA1465120. Variant call files are available on Zenodo (doi: 10.5281/zenodo.20340430). Benchmark variant sets are available from the Genome in a Bottle Consortium (https://ftp-trace.ncbi.nlm.nih.gov/ReferenceSamples/giab//release/AshkenazimTrio/).

## Code availability

All bioinformatics tools used in this manuscript have been mentioned in the text and are freely available on GitHub.

## Author Contributions

The work presented in this manuscript was carried out in collaboration with all authors. All authors made contributions to the conception and design of the work. SG performed DNA extraction and SG and MN performed initial sample quality control. NN, ER and MN performed library preparation and sequencing on the DNBSEQ-T7. AvC, ST and MM performed sequencing quality control and bioinformatic analyses. Variant benchmarking against high-confidence call sets from the Genome in a Bottle Consortium and comparative analyses across library preparation methods were performed by AvC, ST, and MM. AvC performed coverage analyses and Mendelian consistency evaluation for the trio samples. Tables and Figures in the manuscript were generated by AvC and ST. AvC, ST, MM, BG, and NC wrote the manuscript with input from all authors. All authors reviewed and approved the final manuscript. NC, CK, and BG supervised the project.

## Acknowledgements

The authors would like to acknowledge the assistance of DIPLOMICS (www.diplomics.org.za), which is funded by the Department of Science, Technology and Innovation’s South African Research Infrastructure Roadmap, for obtaining some of the GIAB reference materials. This work has benefited from the support and resources provided by the South African Medical Research Council (SAMRC)

